# Genome-first detection of emerging resistance to novel therapeutic agents for SARS-CoV-2

**DOI:** 10.1101/2022.07.14.500063

**Authors:** Manon Ragonnet-Cronin, Rungtiwa Nutalai, Jiandong Huo, Aiste Dijokaite-Guraliuc, Raksha Das, Aekkachai Tuekprakhon, Piyada Supasa, Chang Liu, Muneeswaran Selvaraj, Natalie Groves, Hassan Hartman, Nicholas Ellaby, J. Mark Sutton, Mohammad W. Bahar, Daming Zhou, Elizabeth Fry, Jingshan Ren, Colin Brown, Paul Klenerman, Susanna J. Dunachie, Juthathip Mongkolsapaya, Susan Hopkins, Meera Chand, David I. Stuart, Gavin R. Screaton, Sakib Rokadiya

**Author notes:** Contributed equally (Manon Ragonnet-Cronin, Rungtiwa Nutalai, Jiandong Huo, Aiste Dijokaite-Guraliuc, Gavin R. Screaton, Sakib Rokadiya).

## Abstract

Some COVID-19 patients are unable to clear their infection or are at risk of severe disease, requiring treatment with neutralising monoclonal antibodies (nmAb) and/or antivirals. The rapid roll-out of novel therapeutics means there is limited understanding of the likely genetic barrier to drug resistance. Unprecedented genomic surveillance of SARS-CoV-2 in the UK has enabled a genome-first approach to the detection of emerging drug resistance. Here we report the accrual of mutations in Delta and Omicron cases treated with casirivimab+imdevimab and sotrovimab respectively. Mutations occur within the epitopes of the respective nmAbs. For casirivimab+imdevimab these are present on contiguous raw reads, simultaneously affecting both components. Using surface plasmon resonance and pseudoviral neutralisation assays we demonstrate these mutations reduce or completely abrogate antibody affinity and neutralising activity, suggesting they are driven by immune evasion. In addition, we show that some mutations also reduce the neutralising activity of vaccine-induced serum.

## Introduction

Cases of SARS-CoV-2 infection were first reported in late-December 2019 in Wuhan (Zhou et al., 2020b), and the virus rapidly caused a global pandemic of coronavirus disease 2019 (COVID-19). As of June 2022, over half a billion cases have been reported, with more than 6 million deaths (https://covid19.who.int/). Being a positive-strand RNA virus, although its polymerase has some proofreading ability, SARS-CoV-2 has evolved rapidly with thousands of mutations identified already (Obermeyer et al., 2022). Certain mutations can confer fitness advantages by increasing transmissibility or enabling evasion of humoral responses induced by natural infection or vaccination.

Since the outbreak started several variants of concern (VoC) (https://www.cdc.gov/coronavirus/2019-ncov/variants/variant-classifications.html) have emerged as dominant strains either globally (Dejnirattisai et al., 2022; Liu et al., 2021; Supasa et al., 2021) or regionally (Dejnirattisai et al., 2021b; Zhou et al., 2021). These variants contain multiple mutations mainly found in the gene encoding the viral Spike (S), the major surface glycoprotein crucial for viral infection. The receptor-binding domain (RBD) of the Spike, which initiates viral entry into the host cell by interacting with the host ACE2 receptor, is the major target for potent neutralising antibodies (nmAbs). nmAbs target the RBD in two different ways: most bind to a region on or in close proximity to the ACE2 binding surface of the RBD, whereby they prevent interaction of S with ACE2 and hence block infection (Dejnirattisai et al., 2021a; Yuan et al., 2020a), others bind to non-ACE2 blocking sites on the RBD, and these nmAbs may function to destabilize the trimeric S (Huo et al., 2020; Yuan et al., 2020b; Zhou et al., 2020a).

Drug treatment can drive the evolution of pathogens, leading to rapid selection of advantageous mutations and emergence of resistant strains (Feder et al., 2021). This process can result in failure of treatment; and the spread of resistance may cause new waves of infections. nMAbs are usually prescribed in vulnerable populations where infections persist due to host immunosuppression, further increasing the likelihood of emergence of resistance. (https://assets.publishing.service.gov.uk/government/uploads/system/uploads/attachment_data/file/1039516/S1430_NERVTAG_Antiviral_drug_resistance_and_use_of_Direct_Acting_Antiviral_Drugs_.pdf). There are potentially two ways to avoid mutational escape. Firstly, a cocktail of therapeutics may be developed to simultaneously bind different sites on the target, meaning that to escape, the pathogen will need to evolve two or more mutations, dramatically reducing the chances of escape. Drug cocktails are used to prevent the generation of escape mutations by a number of pathogens such as HIV (Arts and Hazuda, 2012) and TB (Diallo et al., 2021). REGEN-COV is a cocktail of two fully human non-competing nmAbs, casirivimab (REGN10933) and imdevimab (REGN10987), both of which target the ACE2-binding interface of SARS-CoV-2 RBD and function to block RBD/ACE2 interaction (Hansen et al., 2020). *In vitro* experiments demonstrated that the cocktail could neutralise mutants selected (Al-Obaidi et al., 2022) (https://www.covid19treatmentguidelines.nih.gov/therapies/anti-sars-cov-2-antibody-products/anti-sars-cov-2-monoclonal-antibodies/) using single components (Baum et al., 2020; Liu et al., 2021). A previous report also suggested that treatment with REGEN-COV would not lead to the emergence of escape mutants in both preclinical and human studies (Copin et al., 2021).

A second therapeutic strategy to prevent the accrual of escape mutations would be to develop therapeutics to target a conserved epitope that is mutationally constrained, i.e. a mutation of such an epitope would come at a high fitness cost to the pathogen, abrogating any selection advantage. Sotrovimab (VIR-7831 / S309) binds in the region of the N-linked glycan at position 343 of the SARS-CoV-2 RBD; though not interfering with ACE2 binding, it is able to effectively neutralise the virus (Pinto et al., 2020). As this epitope is well conserved among human and animal isolates of clade 1, 2 and 3 Sarbecoviruses (including SARS-CoV-1), sotrovimab (developed from a mAb isolated from a SARS-CoV-1 infected case) was considered to be a broad neutraliser and perhaps able to resist mutational escape even as a monotherapy. It shows an approximately 6-fold reduction in neutralisation of the Omicron variant (Dejnirattisai et al., 2022).

Unprecedented genomic surveillance of SARS-CoV-2 in the UK has enabled a genomic approach to the detection of emerging drug resistance. Here, we report the detection of viral mutations that are associated with drug resistance in patients treated with REGEN-COV (for infection with Delta variant) and sotrovimab (for infection with Omicron variants). We evaluated the binding behaviour of these mutants using surface plasmon resonance (SPR), and examined their impact on the neutralising activity of therapeutic antibodies using pseudoviral assays. Strikingly, the Delta variant was found to acquire mutations at two distinct sites targeted by casirivimab and imdevimab respectively, resulting in severe impairment of neutralising activity of the cocktail. In addition, the Omicron BA.1 variant was found to gain single mutations at multiples sites which completely abolished the binding and neutralisation activity of sotrovimab. Finally, the neutralisation titre of vaccine sera against these escape mutants was significantly reduced compared to the originating strain (i.e. Delta or BA.1 variants).

## Results

### Study population

The present analysis includes all patients who had received treatment in the UK, for whom at least one sample had been collected by 12 April 2022 and for whom a viral genetic sequence was available. Our analysis comprised 21,312 patient sequences sampled before treatment. In the main analysis, sequences were considered post-treatment if patients were sampled at least 10 days after the day of treatment: 1,653 patients treated with one of casirivimab+imdevimab, molnupiravir, pnirmatrelvir plus ritonavir (Paxlovid), remdesivir or sotrovimab.

### Post-treatment mutation analysis

We compared amino acid frequencies between pre- and post-treatment sequences. stratifying analyses by treatment, variant (Delta, BA.1 or BA.2), and gene. Nine amino acid residues displayed a significant (p<0.001) frequency change in post-treatment sequences compared to pre-treatment sequences, suggesting possible evidence of selection. All treatment-emergent substitutions were in the Spike RBD region: E406D/Q, G446S/V, Y453F and L455F/S in patients infected with Delta and treated with casirivimab+imdevimab; P337R/S and E340A/D/K/V, K356T and R493Q in patients infected with BA.1 and treated with sotrovimab; and E340K in patients infected with BA.2 and treated with sotrovimab (**Figure 1**). For molnupiravir, remdesivir and paxlovid, no significant (p<0.001) mutations were observed in the available data.

**Figure 1.**
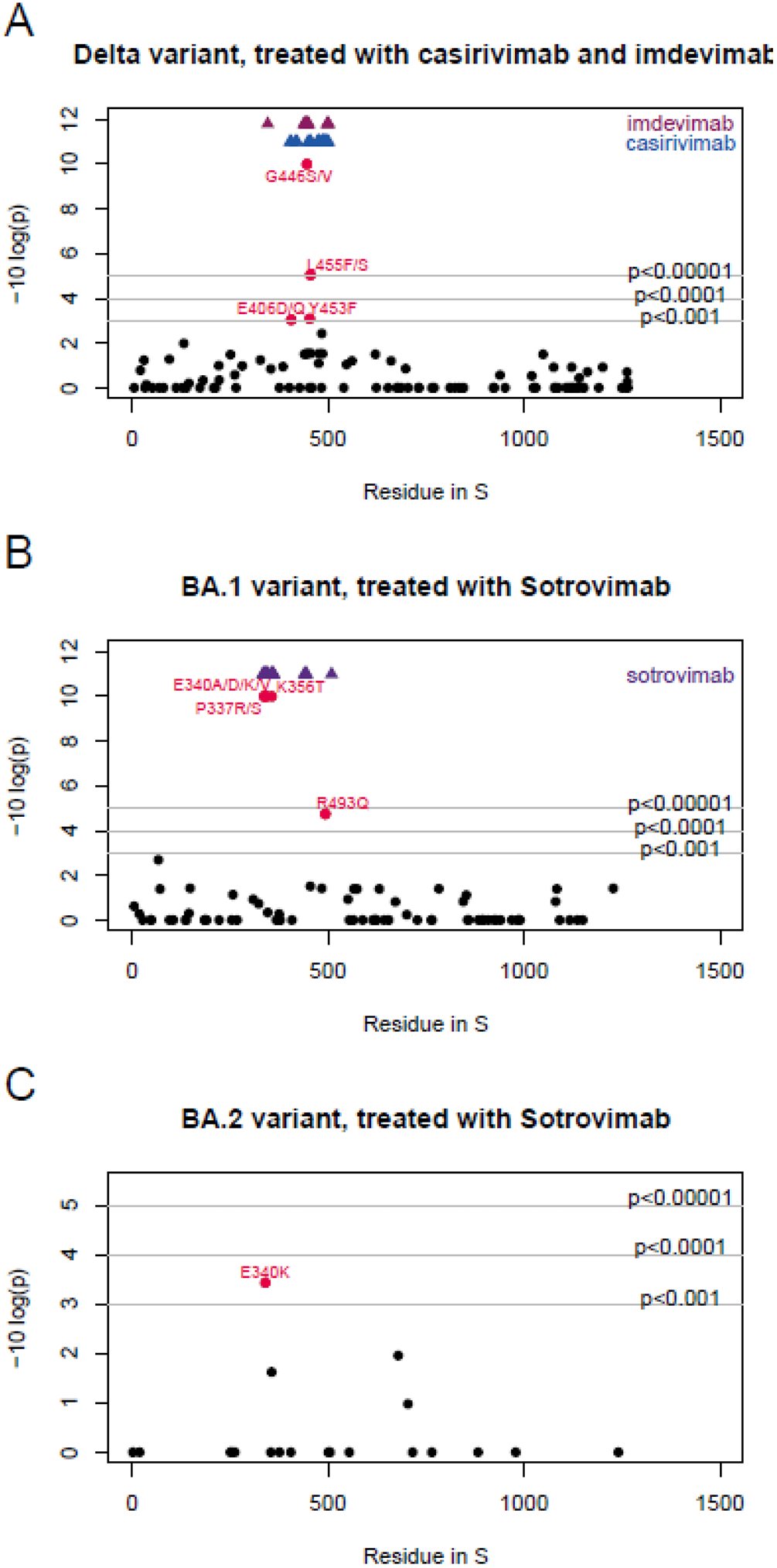
P-values for differences in spike amino acid frequencies between pre- and post-treatment sequences. (A) patients infected with Delta and treated with casirivimab/imdevimab, (B) patients infected with BA.2 and treated with sotrovimab, and (C) patients infected with BA.2 and treated with sotrovimab. Amino acid frequencies were compared between pre-and post-treatment samples (at least 10 days after treatment) at each site in the spike sequence alignment. P-values for each site were calculated using a Fisher’s test, and p-values were log-transformed and inversed for visualisation so that sites with diverging values appear higher up on the figure. Only sites with some variability (>1 amino acid) are shown. The horizontal lines indicate p-value thresholds of p<0.001, p<0.0001 etc. Residues with diverging frequencies (p<0.001) are highlighted in red, with the observed amino acid change indicated in text. Residues known to interact with each drug are indicated in blue and purple at the top of the figure. The numbers differ slightly from those in **Table 1** because not all gene regions were sufficiently high quality for downstream sequence analysis. See also **Figure S1**.

Restricting the calculation to the three groups with identified associations: patients infected with Delta and treated with casirivimab+imdevimab and patients infected with BA.1 or BA.2 and treated with sotrovimab (**Table 1**), a total of 86/959 (8.97%) post-treatment (≥1 day) patients had at least one of the identified mutations, compared to 16/7,788 (0.20%) pre-treatment patients (**Table 2**; p<10^−16^). Eleven post-treatment patients had >1 mutation: three patients infected with Delta treated with casirivimab+imdevimab had a combination of G446V and L455F, one had G446S and L455 and one had G446V and Y453F. We examined the raw reads and confirmed that for all of these patients, both mutations were present on most contiguous raw reads. Among BA.1 patients treated with sotrovimab, four had a combination of E340A and R493Q, one had E340D and R493Q and one had K356T and R493Q.

**Table 1.** Data set sizes. Note that some patients received multiple courses of treatment and thus may be counted more than once in the table.

| Treatment | Variant | Number of patients (pre-treatment) | Number of patients (≥ 1 day post-treatment) | Number of patients (≥ 5 day post-treatment) | Number of patients (≥ 10 day post-treatment) | Number of patients (≥ 14 day post-treatment) |
| --- | --- | --- | --- | --- | --- | --- |
| Casirivimab and imdevimab | BA.1 | 137 | 85 | 73 | 64 | 58 |
| Casirivimab and imdevimab | BA.2 | 0 | 11 | 12 | 12 | 12 |
| Casirivimab and imdevimab | delta | 1557 | 227 | 123 | 67 | 50 |
| Molnupiravir | BA.1 | 1411 | 150 | 104 | 67 | 41 |
| Molnupiravir | BA.2 | 228 | 17 | 11 | 8 | 5 |
| Molnupiravir | delta | 24 | 7 | 6 | 4 | 1 |
| Paxlovid | BA.1 | 276 | 18 | 8 | 6 | 2 |
| Paxlovid | BA.2 | 598 | 40 | 15 | 10 | 5 |
| Paxlovid | delta | 0 | 1 | 1 | 1 | 1 |
| Remdesivir | BA.1 | 872 | 397 | 305 | 258 | 227 |
| Remdesivir | BA.2 | 187 | 92 | 76 | 65 | 65 |
| Remdesivir | delta | 3054 | 703 | 334 | 201 | 133 |
| Sotrovimab | BA.1 | 3221 | 380 | 240 | 148 | 114 |
| Sotrovimab | BA.2 | 1338 | 112 | 50 | 25 | 18 |
| Sotrovimab | delta | 24 | 5 | 5 | 2 | 2 |

**Table 2.** Frequency of each mutation.

| Variant | treatment | gene | mutation | frequency<br>in pre-treatment<br>patients | Frequency<br>in treated<br>patients | p value |
| --- | --- | --- | --- | --- | --- | --- |
| Delta | Casirivimab<br>and<br>imdevimab | spike | E406D/Q | 0 (0%) | 1 (1.2%) | p<10-3 |
| Delta | Casirivimab<br>and<br>imdevimab | spike | G446S/V | 2 (0.1%) | 8 (9.8%) | p<10-16 |
| Delta | Casirivimab<br>and<br>imdevimab | spike | Y453F | 0 (0%) | 2 (2.4%) | p<10-3 |
| Delta | Casirivimab<br>and<br>imdevimab | spike | L455F/S | 2 (0.1%) | 4 (4.9%) | p<10-5 |
| BA.1 | sotrovimab | spike | P337R/S | 0 (0%) | 12 (5.7%) | p<10-17 |
| BA.1 | sotrovimab | spike | E340A/D/K/V | 4 (0.1%) | 31 (14.7%) | p<10-18 |
| BA.1 | sotrovimab | spike | K356T | 0 (0%) | 5 (2.4%) | p<10-19 |
| BA.1 | sotrovimab | spike | R493Q | 1 (0.03%) | 4 (1.9%) | p<10-4 |
| BA.2 | sotrovimab | spike | E340K | 1 (0.05%) | 2 (8%) | p<10-4 |

To determine sensitivity, the analysis was repeated with a different threshold for post-treatment sequences: at least one or five days after treatment. While datasets were much larger when a shorter interval was considered, the strength of the signal became stronger as the interval was lengthened, lending support to the validity of our findings (**Figure S1**).

### Frequency of mutations in UK genomic database

For each mutation identified, we ascertained its frequency in the UK genomic database from September 2022 onwards. The frequency of the mutations listed previously within the UK genomic data set for mutations associated with casirivimab and imdevimab in Delta sequences (n=752,585) were: 6 E406D; 7 E406Q; 163 G446S; 1,946 G446V, 12 Y453F; 179 L455F; 1 L455S. The frequency of mutations post-sotrovimab treatment with the BA.1 variant (n=702,940) was: 11 P337R; 32 P337S; 39 E340A; 82 E340D; 52 E340K; 5 E340V; 57 K356T; 1214 R493Q. The frequency of mutations post-sotrovimab treatment with the BA.2 variant (n=407,161) was: 10 E340K. As above, a total of 86/959 (8.97%) post-treatment (≥1 day) patients had at least one of the identified mutations; in contrast, the frequency of any mutation in the variants of interest in the genomic surveillance dataset was 3,653/1,862,686 (0.20% identical to the frequency in the pre-treatment dataset; p<10^−16^).

Overall, these data demonstrate a significant enrichment of mutations in the post-treatment sequences compared to the pre-treatment group and compared to the genomic database as a whole, strongly implicating them as mutations selected for escape from nmAb therapy.

### Mapping of mutations to the Spike

Figure 2. shows the positions of the mutations found to be of high significance. All mutations occur in the RBD of the SARS-CoV-2 Spike.

**Figure 2.**
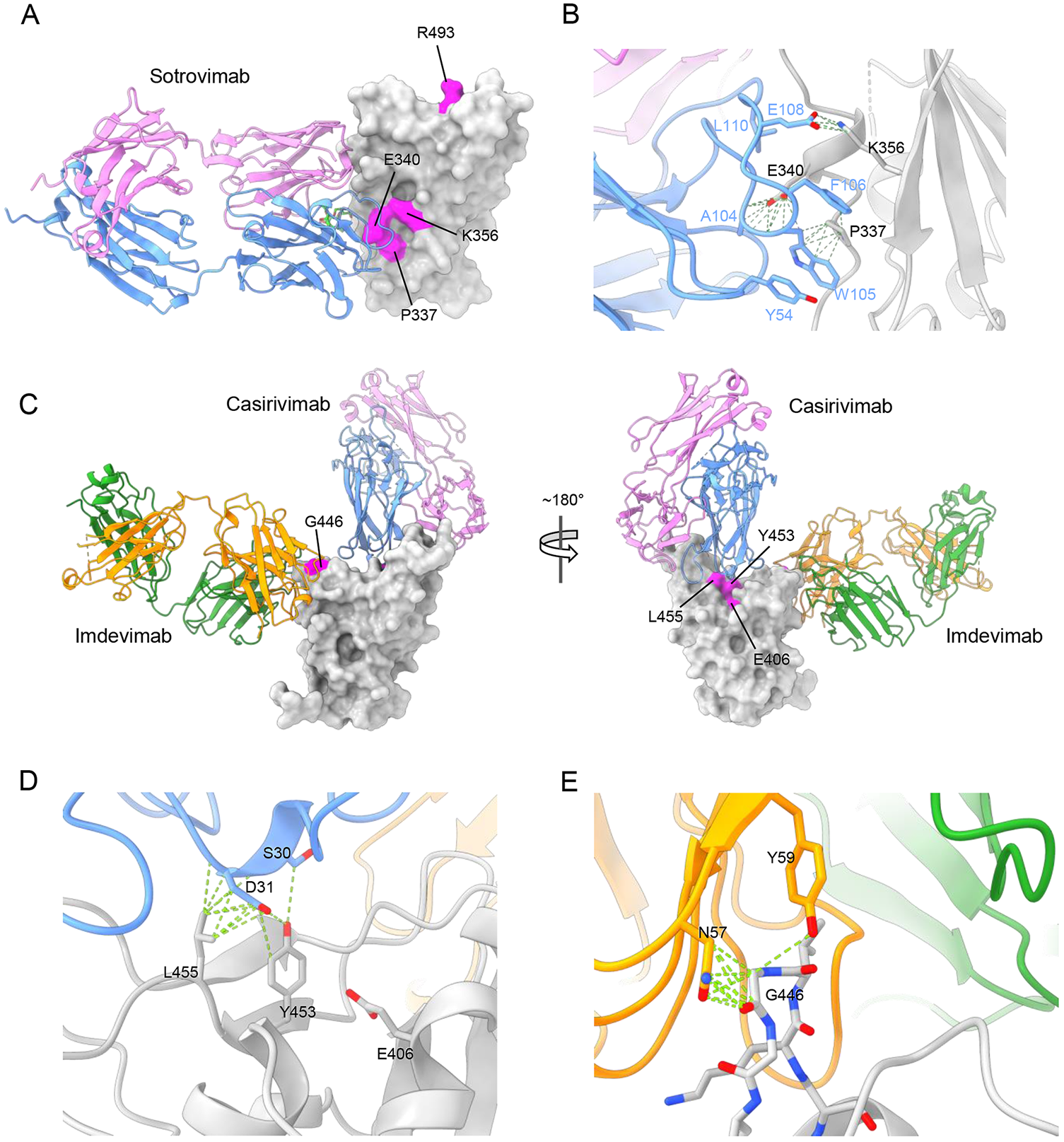
Structural modelling of mutations mapped to the Spike RBD. (A) Model of the Omicron RBD (PDB: 7TLY) docked with S309 (sotrovimab). Omicron RBD is shown as a grey surface from an approximate front view, S309 as cartoon ribbons with heavy and light chains coloured separately. Mutation sites mapped to the RBD surface are coloured magenta and labelled. (B) Close-up view of the interface between the P337, E340, K356 patch of residues with the S309 heavy chain. Potential hydrogen bonds and hydrophobic interactions are shown as green dashed lines. (C) Model of the Delta RBD docked with REGEN-COV nmAbs casirivimab and imdevimab shown from approximate front (left) and back (right) views. Delta RBD is shown as a grey surface and mutation sites E406, G446, Y453 and L455 are coloured magenta and labelled. (D) Close-up view of the interface between E406, Y453 and L455 with casirivimab. (E) Close-up view of the interface between G446 with imdevimab. Potential hydrogen bonds and hydrophobic interactions are shown as green dashed lines.

In **Figure 2A** the mutations associated with sotrovimab treatment are mapped to the structure of the Omicron BA.1 RBD and sotrovimab complex (PDB: 7TLY) (McCallum et al., 2022a). Note that RBD residues 337, 340 and 356 cluster tightly forming an interaction hotspot with the antibody heavy chain CDR3 in particular (**Figure 2B)**. W105 and F106 of the CDR3 form a key 4-layer hydrophobic sandwich with residues 337 and 356 of the RBD (W105:P337:F106:K356), whilst E340 pins down the CDR3 loop by a remarkable set of interactions with the amide nitrogens of residues 104-106, which are arranged rather as an open helix capped with exquisite specificity by E340. This suggests that the observed mutations P337R/S; E340A/D/K/V; K356T will all disrupt this binding hotspot. In contrast Q493R is distal to the epitope, on the edge of the ACE2 footprint (**Figure 2A**), so there is no obvious reason for this mutation to affect antibody binding.

The mutations associated with casirivimab and imdevimab treatments are shown in **Figure 2C**, mapped to the structure of the Delta RBD containing the L452R mutation (PDB:7ORB) (Liu et al., 2021), where the binding of casirivimab and imdevimab is inferred from the reported structure of the complex with early pandemic RBD (PDB:6XDG, the RMSD in C**α** positions between early pandemic and Omicron BA.1 RBDs is 1.16 Å and we are confident that this inference is secure). The mutations observed fall into two areas on the surface of the RBD. Positions 406, 453 and 455 are clustered together at the back of the neck region (Dejnirattisai et al., 2021) lying under the CDR1 of the casirivimab heavy chain and forming a nest of interactions (**Figure 2D**). These mutations would be expected to affect binding of this antibody. In contrast G446 rests tightly against N57 and Y59 of the light chain CDR2 of imdevimab (**Figure 2E**) and any change to a larger side chain such as the G446S/V mutations observed, would be expected to abrogate binding.

### Experimental measurement of escape by mutants identified from patients treated with REGEN-COV

We constructed a panel of pseudotyped lentiviruses (Di Genova et al., 2020) expressing the Spike from the identified escape mutants (**Figure 3**). Pseudoviral neutralisation assays showed that activity of imdevimab against the Delta+G446V mutant was completely knocked out, whilst casirivimab showed >10-fold reductions in the neutralization titre of Delta+Y453F (16-fold), Delta+L455F (17-fold) and Delta+L455S (155-fold), compared to the wild-type Delta variant **(Figure 3A,C)**

**Figure 3.**
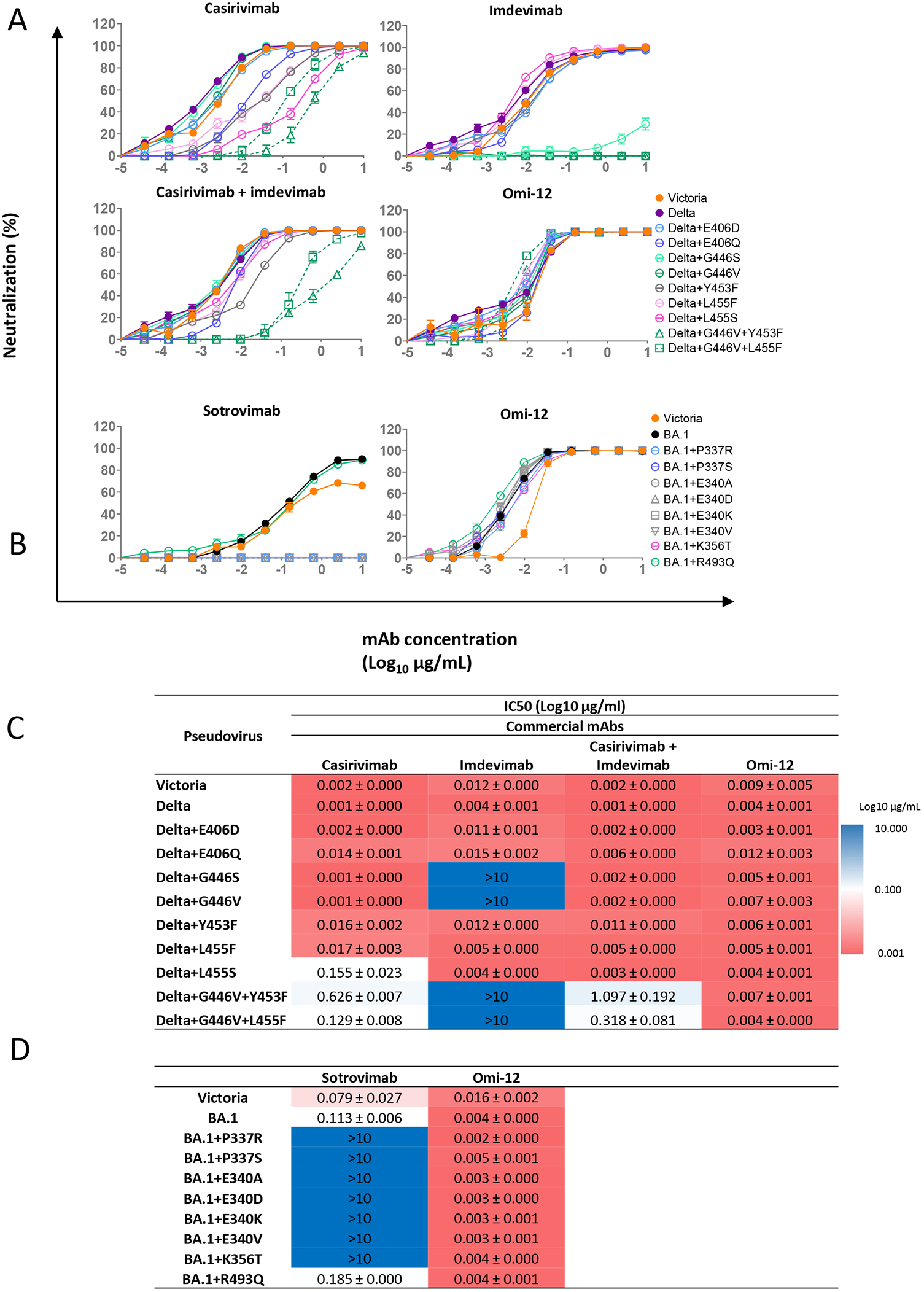
Neutralization escape caused by RBD mutations. (A) Pseudoviral neutralization curves of the indicated Delta variants with REGEN-COV nmAbs. Comparison is made with Omi-12 A VH1-58 mAb which is not sensitive to the mutations found following REGEN-COV treatment. (B) Pseudovirus neutralization curves for BA.1 sotrovimab mutants. (C, D) Neutralization IC50 titres for neutralizations shown in A, B.

As casirivimab remained fully active against the Delta+G446V mutant, and imdevimab was still able to potently neutralize the Delta+Y453F and Delta+L455F/S mutants, the combination of casirivimab and imdevimab retained neutralization potency against all these single mutants. However, the combined mutations of Delta+G446V+Y453F and Delta+G446V+L455F not only led to complete knock-out of the neutralising activity of imdevimab, but also severe knock-down of casirivimab activity. As a result, the neutralisation titre of casirivimab+imdevimab was reduced 1097-fold against Delta+G446V+Y453F and 318-fold against the Delta+G446V+L455F. This is consistent with the finding of these pairs of mutations occurring together on single Delta RBD sequences described above.

To confirm that the observed effects on neutralization were directly attributable to the change in RBD/nmAb interaction, we measured the affinity of nmAbs and RBD mutants by surface plasmon resonance (SPR) (**Figure S2**,**S3, Table 3A**). This analysis also showed that the G446V mutation almost abolished the binding of imdevimab, and in the meantime caused a modest reduction (1.8-fold) in the binding affinity of casirivimab (**Table 3A)**. The L455S, E406D and E406Q single mutations mainly affect casirivimab. SPR analysis showed a 369-fold, 20-fold and 38-fold decrease in the affinity of casirivimab for Delta+L455S, Delta+E406D and Delta+E406Q respectively. The neutralisation titre of casirivimab was reduced 65-fold, 2-fold and 12-fold against these three mutants respectively (**Figure 3C, Table 3A**). However, since imdevimab was unaffected, the casirivimab+imdevimab combination retained potent neutralising activity against these mutants.

**Table 3.** SPR analysis of RBD/nmAb interaction KD and fold change **A, B** Summary of binding affinity between RBDs and therapeutic mAbs (A) casirivimab+imdevimab, (B) sotrovimab). The fold reduction in affinity between RBD mutants and wild-type RBD is calculated. The number labelled with a star indicates a fold increase in affinity. See also **Figures S2** and **3**.

| RBD | Casirivimab |  | Imdevimab |  |
| --- | --- | --- | --- | --- |
|  | K<sub>D</sub> (nM) | Fold reduction | K<sub>D</sub> (nM) | Fold reduction |
| Delta RBD WT | 0.36 | - | 9.4 | - |
| Delta RBD+E406D | 7.1 | 20 | 15 | 1.6 |
| Delta RBD+E406Q | 14 | 38 | 8.9 | 1.1* |
| Delta RBD+G446S | 0.56 | 1.6 | 734 | 78 |
| Delta RBD+G446V | 0.64 | 1.8 | Very weak binding |  |
| Delta RBD+Y453F | 67 | 186 | 9.8 | 1.0 |
| Delta RBD+L455F | 44 | 122 | 11 | 1.2 |
| Delta RBD+L455S | 133 | 369 | 9.1 | 1.0 |
| Delta RBD+G446V+Y453F | 125 | 347 | Very weak binding |  |
| Delta RBD+G446V+L455F | 69 | 192 | Very weak binding |  |

**B**
| RBD | Sotrovimab |  |
| --- | --- | --- |
|  | K<sub>D</sub> (nM) | Fold reduction |
| BA.1 RBD WT | 0.17 | - |
| BA.1 RBD+P337R | 753 | 4428 |
| BA.1 RBD+P337S | 332 | 1951 |
| BA.1 RBD+E340A | 3415 | 20088 |
| BA.1 RBD+E340D | 764 | 4494 |
| BA.1 RBD+E340K | 3441 | 20241 |
| BA.1 RBD+E340V | 345 | 2027 |

Interestingly, an additive effect on reducing casirivimab binding was seen for the combination of mutations resulting in an overall 347-fold and decrease in affinity for G446V+Y453F and 192-fold decrease for G446V+L455F. As expected, binding of imdevimab to Delta+G446V+Y453F and Delta+G446V+L455F was almost completely impaired. Overall, the acquisition of double mutations has rendered substantial loss in sensitivity to the REGEN-COV regime.

### Experimental measurement of escape by mutants identified from patients treated with sotrovimab

BA.1 mutations P337R/S and E340A/D/K/V, led to complete knock out of neutralisation by sotrovimab (**Figure 3B,D**). Although the BA.1+R493Q (reversion to Wuhan wild type) was also identified as a post-treatment emergent mutation, no obvious effect on the neutralising activity of sotrovimab was observed. The RBDs of BA.1+P337R/S and BA.1+E340A/D/K/V were successfully expressed to allow examination of their binding with sotrovimab (**Figure S3**). The affinity of sotrovimab was reduced by 1951-fold to 20241-fold compared to the wild-type BA.1 RBD, explaining why these mutants were resistant to sotrovimab neutralization (**Table 3D**).

### Neutralization of escape mutants by vaccine serum

Neutralization assays were performed using serum obtained 28 days following a third dose of Pfizer-BioNtech vaccine BNT162b2 (Cele et al., 2021) **(Figure 4)**. Following 3 doses of BNT162B a 1.9-fold and 1.5-fold decrease was observed for Delta+G446V+Y453F and Delta+G446V+L455F respectively, compared to wild-type Delta (p<0.0001); whilst a 2-fold, 1.2-fold and 3.8-fold reduction was seen for BA.1+P337S, BA.1+E340K and BA.1+K356T respectively compared to wild-type BA.1 (p<0.0001, p=0.0082 and p<0.0001).

**Figure 4.**
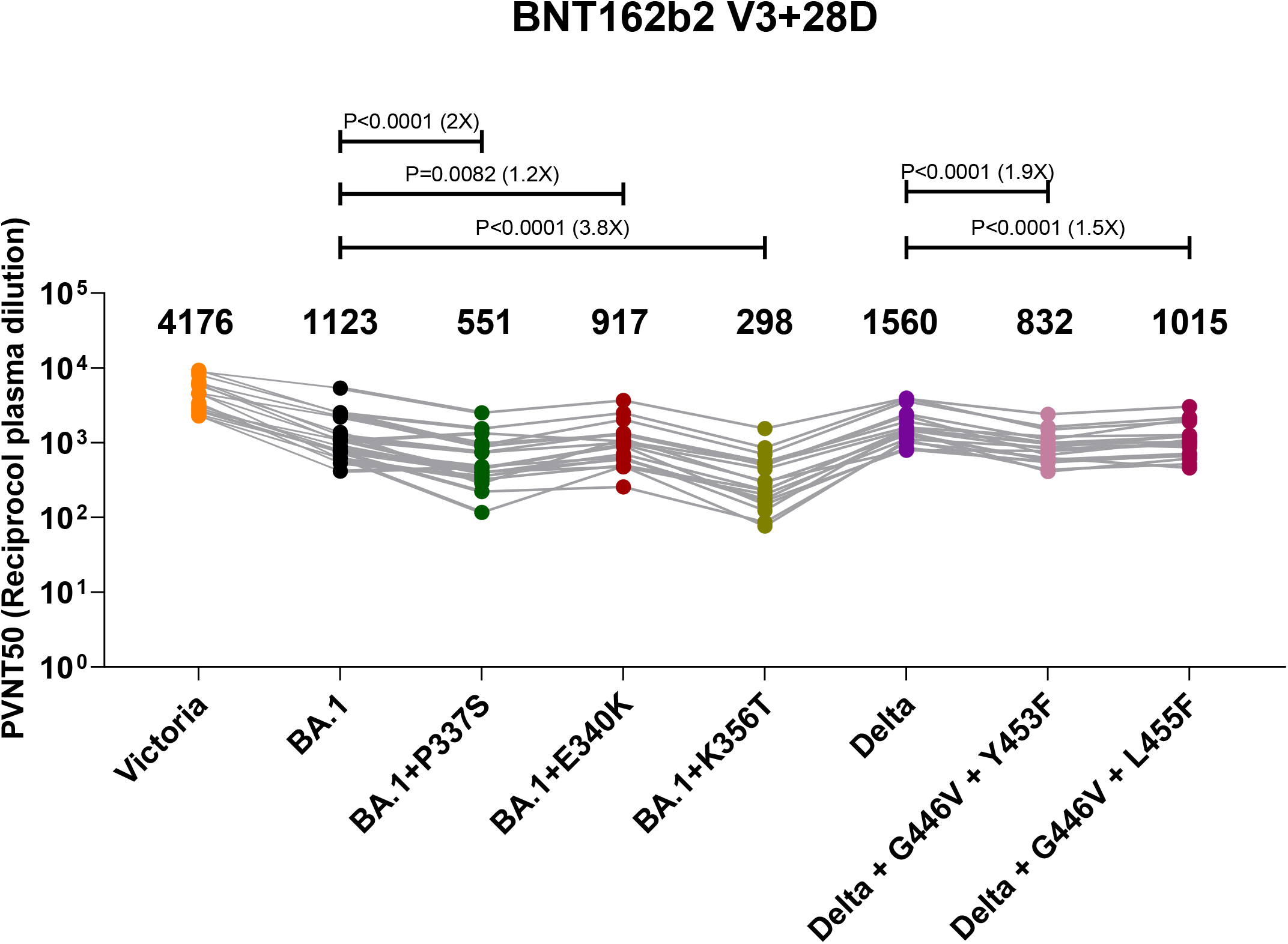
Pseudoviral neutralization IC50 titres of third dose Pfizer BioNTech vaccine serum. IC50 titres for Delta REGEN-COV induced mutations and BA.1 sotrovimab induced mutations are compared with titres for ancestral strain Victoria, Delta and BA.1. Geometric mean titres are shown above each column. The Wilcoxon matched-pairs signed rank test was used for the analysis and two-tailed P values were calculated.

## Discussion

Individuals infected with the currently dominant SARS-CoV-2 Omicron variant have been shown to have a lower likelihood of severe disease and hospitalisation compared with previous variants. However, a large number of people still suffer from severe disease (Wolter et al., 2022) and this proportion could be higher in populations with lower levels of infection- or vaccine-induced immunity.

Although the current mortality rates are much lower than in 2020 (https://www.ons.gov.uk/peoplepopulationandcommunity/healthandsocialcare/conditionsanddiseases/articles/coronaviruscovid19latestinsights/deaths), as of June 2022 over 300 people died from COVID-19 every week within the UK (https://coronavirus.data.gov.uk/details/deaths). Individuals who are unable to mount an adequate immune response from vaccination or for whom vaccination is not recommended are at particular risk. It is this vulnerable population, who tend to suffer from chronic COVID-19 infections, who are targeted to receive nmAb therapies either therapeutically or prophylactically. In the UK, the highest-risk clinical subgroups who are immunosuppressed are eligible for these therapies (https://www.gov.uk/government/publications/higher-risk-patients-eligible-for-covid-19-treatments-independent-advisory-group-report/defining-the-highest-risk-clinical-subgroups-upon-community-infection-with-sars-cov-2-when-considering-the-use-of-neutralising-monoclonal-antibodies).

While commercial anti-SARS-CoV-2 therapeutic mAbs have been shown to be effective treatments for COVID-19 (Gupta et al., 2021; Weinreich et al., 2021), various studies have reported severe reductions or complete knock-out of their neutralising activities against Omicron variants (Dejnirattisai et al., 2022; McCallum et al., 2022b; Nutalai et al., 2022). As sotrovimab was shown to be unable to effectively neutralise Omicron BA.2, in April 2022 the FDA announced that sotrovimab was no longer authorized to treat COVID-19 as BA.2 became the dominant variant (https://www.fda.gov/drugs/drug-safety-and-availability/fda-updates-sotrovimab-emergency-use-authorization). However, in the UK, sotrovimab remains in clinical use.

In a recent study the Delta variant was reported to develop P337L/T and E340K/A/V resistance mutations in patients treated with sotrovimab (Rockett et al., 2022). Here we report the identification of BA.1 escape mutations in patients who received sotrovimab treatment. In addition to mutations occurring at the P337 and E340 residues, we also identify a novel K356T mutation. These mutations abolish the binding and hence neutralising activity of sotrovimab. Q493R is also found, a reversion to the sequence found in early pandemic viruses and in BA.4/5. This mutation is distal to sotrovimab footprint, has no effect on antibody binding but has been reported to increase the affinity for ACE2 (Wang et al., 2022), suggesting improved receptor binding rather than escape from antibody binding may be the driver for selection. These observations suggest that monotherapy is likely to be impacted by emerging variants and induce treatment-emergent resistance, even if the drug targets an epitope that is well conserved among Sarbecoviruses.

In contrast to the single agent sotrovimab, the REGEN-COV regime, containing a combination of two nmAbs that target non-overlapping epitopes, would be expected to be more resistant to mutational escape. Indeed, previous studies have shown that REGEN-COV was able to effectively prevent emergence of escape mutants not only *in vitro*, but also in *in vivo* animal and human studies (Baum et al., 2020; Copin et al., 2021). However, in this detailed study, we observe that treatment with the dual agent REGEN-COV led, in some individuals, to the Delta variant acquiring pairs of mutations that simultaneously impair the binding of both components of REGEN-COV, leading to up to 1000-fold reduction in neutralization titres. All the mutations we identified had been predicted in a mapping exercise where the impact of every potential mutation in the spike protein was tested. The study revealed that pseudoviruses with an E406W mutation were able to escape from both REGEN-COV compounds (Starr et al., 2021). This mutation did not occur in our small dataset. Two nucleotide changes are required for this change in amino acid; however, single nucleotide changes at the site were identified and found to be significant.

It is uncertain how the virus was able to gain the combined resistance mutations during therapy, however, accelerated viral evolution has been documented in immunocompromised patients who could suffer from persistent SARS-CoV-2 infections for many months, with mutations found predominantly in the RBD and other regions of the Spike (Choi et al., 2020). One possibility is that viruses harbouring mutations resistant to one component of REGEN-COV might have already emerged in such patients prior to the cocktail treatment, and the medication then drove selection of a second mutation, leading to an overall impairment of the therapy, perhaps accelerated by viral recombination. If accelerated virus evolution has facilitated escape via a bystander effect (for instance mutations driven by modulation of receptor binding) this might be an additional argument for attempting to find neutralising antibodies that bind in more conserved regions, although we find that increased receptor affinity is selected in some BA.1 infected patients treated with sotrovimab, which binds a conserved. However, viruses bearing single escape mutations were identified in patients under the REGEN-COV treatment. In effective combination therapy, these mutants would be neutralised by one of the components. This raises the question of whether the concentration of the mAbs might be unable to reach the desired level *in vivo*, for example due to limited or differential bioavailability in certain parts of the body, creating a favourable environment for viruses to develop resistance.

The simplest way to mitigate escape is probably to use a more complex cocktail of non-competing mAbs, indeed it has been shown that such a combination was able to retain antiviral potency through up to eleven consecutive serial passages (Copin et al., 2021). Combining mAbs with antivirals is another option, or devising clinical approaches based on patient profile together with using the correct dose for bioavailability. It could also be important to perform genotyping for variants prior to administration of mAb therapy, particularly in chronically infected immunocompromised cases (Greninger et al., 2022). However, patients prescribed treatment for COVID-19 infections are usually started on therapy the same day, and so the turnaround time between sampling and sequence analysis would have to be substantially shortened for clinical use.

Finally, it’s concerning that the neutralisation titre of vaccine serum was reduced against escape mutants evolved from both treatment regimes in two different virus variants. This is not altogether surprising, as the nmAbs chosen for therapeutic use target important neutralising epitopes on the SARS-CoV-2 RBD. Whether nmAb therapy can drive the generation of novel highly transmissible variants is not clear; our study using *in vitro* neutralization gives no indication how fit these variants would be in the general population. It also seems unlikely that mAb-driven escape in the extremely small number of patients given therapy will markedly accelerate generation of novel variants compared to what is happening in the pandemic at large, with millions of infections occurring every day, in an increasingly naturally exposed or vaccinated population, where the selection pressure for antibody escape is already extreme. However, the repeated and perhaps inconsistent use of nmAb therapy in chronically infected individuals, who have been documented to harbour virus for months and in some cases more than a year, should be closely monitored. The analysis of post-treatment sequence datasets and potential transmission of post-treatment emergent mutations is performed regularly by the UK Health Security Agency and published online (https://www.gov.uk/government/publications/covid-19-therapeutic-agents-technical-briefings).

In summary, we demonstrate here mutational changes in viruses isolated from patients treated with nmAbs. The mutational profiles of patients treated with sotrovimab or REGEN-COV are strikingly different and the mutations map to the binding sites for the mAb on Delta or BA.1 RBD. The corresponding mutations impair the binding of nmAbs to Spike RBD, resulting in reduced neutralization titre. Strikingly, for REGEN-COV, viruses evolve pairs of mutations to escape both components of the antibody cocktail.

## Limitations of the study

These studies used *in vitro* neutralization assays and may underestimate the neutralization potential of mAb *in vivo*, where the effects of antibody dependent cell mediated cytotoxicity and complement may increase activity. In addition, using *in vitro* systems we are unable to determine whether the escape mutations selected by nmAb therapy would be fit to compete with natural viral variants in natural infections. We did not look at deep sequence data to look at changes in frequencies of minor variants over time, our sequences are consensus reads. Our single amino acid approach may miss compensatory mutations that do not come out as significant in a large-scale analysis but may be important within patients who have already developed one treatment-emergent substitution. This study did not examine T cells which contribute to the host defence and are less impacted by mutations.

## Acknowledgements

We thank Andrew Balmer for examining the raw reads to confirm linkage between mutations, and the entire Genomics Public Health Analysis team and the COVID-19 therapeutics programme, at UK Health Security Agency. We thank all the teams involved in sampling, sequencing and processing SARS-CoV-2 viruses and sequences. We are grateful to all the volunteers who donated blood for the study.

This work was supported by the Chinese Academy of Medical Sciences (CAMS) Innovation Fund for Medical Science (CIFMS), China (grant number: 2018-I2M-2-002) to D.I.S. and G.R.S. We are also grateful for support from the Red Avenue Foundation and the Oak Foundation. G.R.S. and J.Ren are supported by the Wellcome Trust (101122/Z/13/Z), D.I.S. and E.E.F. by the UKRI MRC (MR/N00065X/1). D.I.S., S.J.D and G.R.S. are Jenner Investigators. P.K. is a NIHR Senior Investigator and P.K. is supported by Wellcome (WT109965MA). S.J.D. is supported by an NIHR Global Research Professorship (NIHR300791)

This work was also supported by the UK Department of Health and Social Care as part of the PITCH (Protective Immunity from T cells to Covid-19 in Health workers) Consortium, and UKRI (MR/W02067X/1), with contributions from UKRI/NIHR through the UK Coronavirus Immunology Consortium (UK-CIC), the Huo Family Foundation and The National Institute for Health Research (UKRIDHSC COVID-19 Rapid Response Rolling Call, Grant Reference Number COV19-RECPLAS).

We thank the staff of the MRC Human Immunology Unit for access to their Biacore Facility.

We acknowledge joint Centre funding from the UK Medical Research Council and Department for International Development (MR/R015600/1). This work is also supported by the National Institute for Health Research Health Protection Research Unit in Modelling Methodology, the Abdul Latif Jameel Foundation and the EDCTP2 programme supported by the European Union.

## Author Information

These authors contributed equally: M.R-C., R.N., J.H., A.D-G.

## Contributions

M.R.C., S.R., G.R.S and D.I.S. conceived of the study. M.R.C. performed the post-treatment analyses. N.G. performed the UK genomic surveillance analyses. R.N., A.D-G., R.D., A.T., P.S., C.L., M.S., D.Z. and J.H. performed experiments, J.H., J.M., M.C. designed experiments, P.K. and S.J.D. oversaw the OPTIC study for the collection of vaccine serum, as part of the PITCH consortium. M.B., J.R., E.F., D.I.S performed structure analyses. N.G., H.H., N.E., M.S., C.B., S.H. and S.R. analysed sequence data. M.R.C., G.R.S., J.H. and D.I.S wrote the first draft of the manuscript. All authors reviewed and approved the final manuscript.

## Figure legends

**Figure S1.**
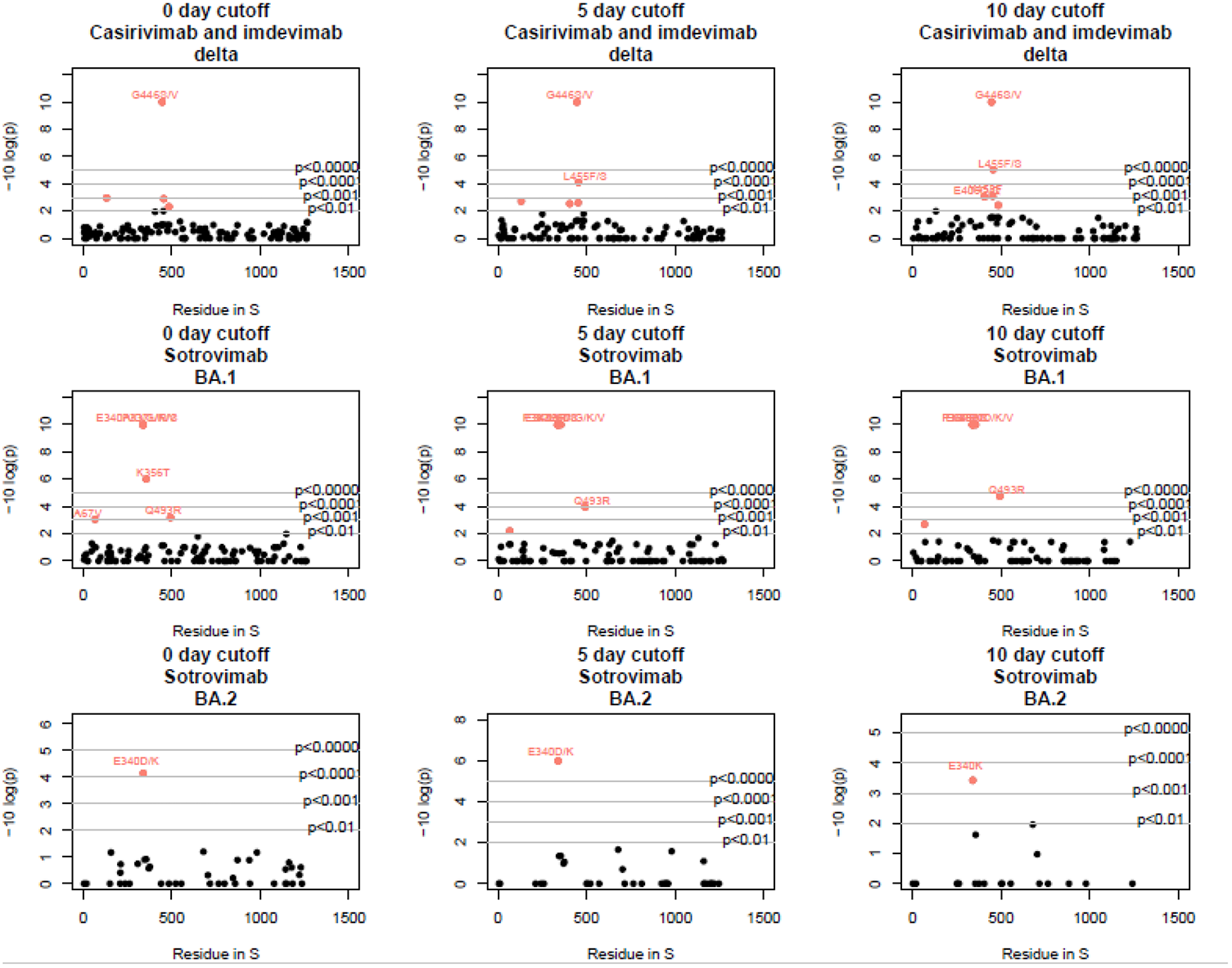
P-values for differences in spike amino acid frequencies between pre- and post-treatment sequences. The indicated cut off dates, following the nmAb treatment were used for the acquisition of the post treatment sample.

**Figure S2.**
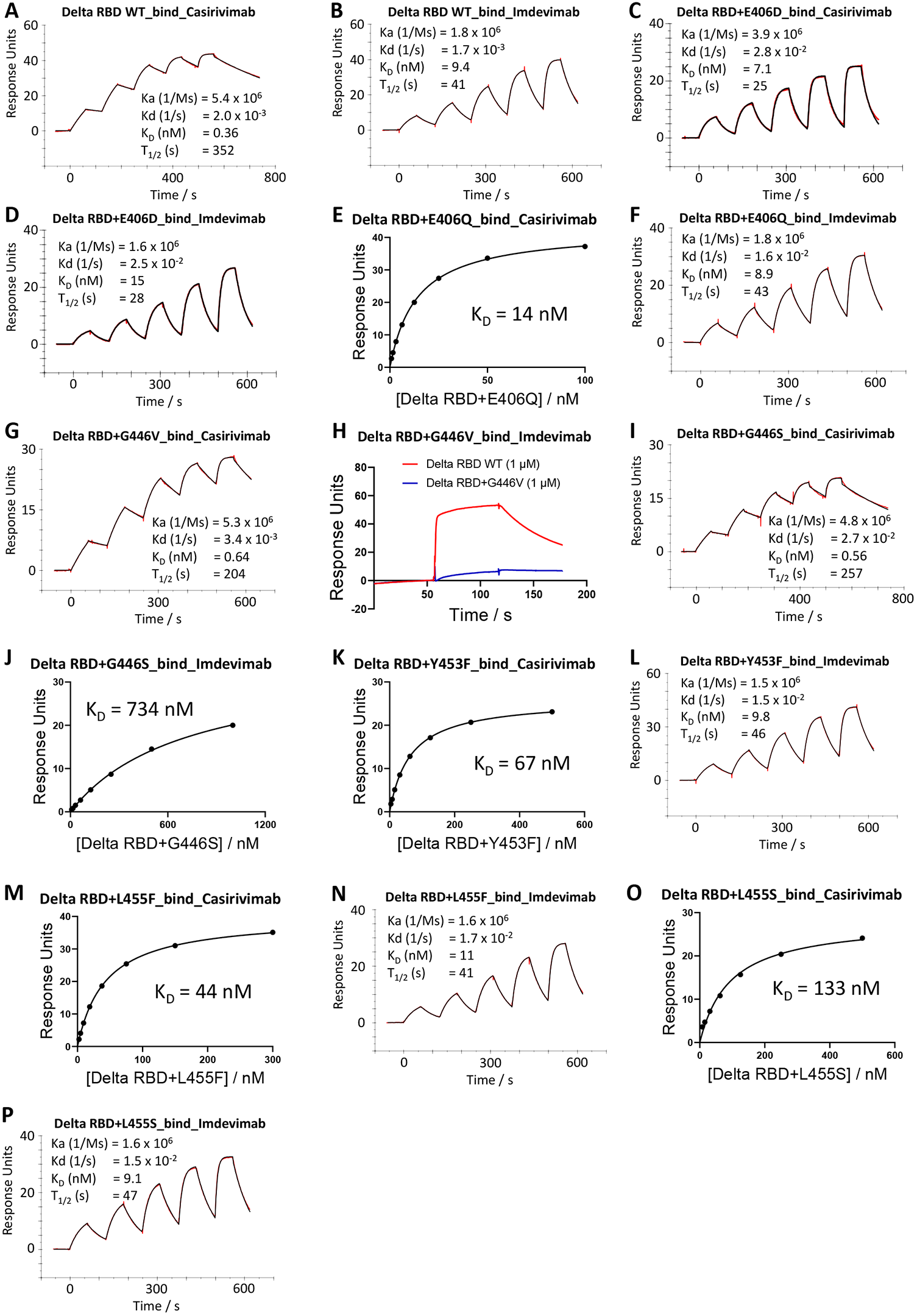
Surface plasmon resonance (SPR) analysis of interaction between Delta and BA.1 RBD mutants and therapeutic mAbs. (A-D; F-G; I, L, N, P) Sensorgrams showing the binding of wild-type Delta RBD and Delta RBD mutants to casirivimab/imdevimab, with affinity and kinetic parameters shown. (E, J, K, M, O) 1:1 binding equilibrium analysis of binding of Delta RBD mutants to casirivimab/imdevimab, with affinity values shown. (H) Binding of Delta RBD+G446V to imdevimab is severely reduced compared to that of wild-type Delta RBD, so that the binding could not be accurately determined, as shown by a single-injection of 1 μM RBD over sample flow cells containing imdevimab. Related to **Figure S3** and **Table 3**.

**Figure S3.**
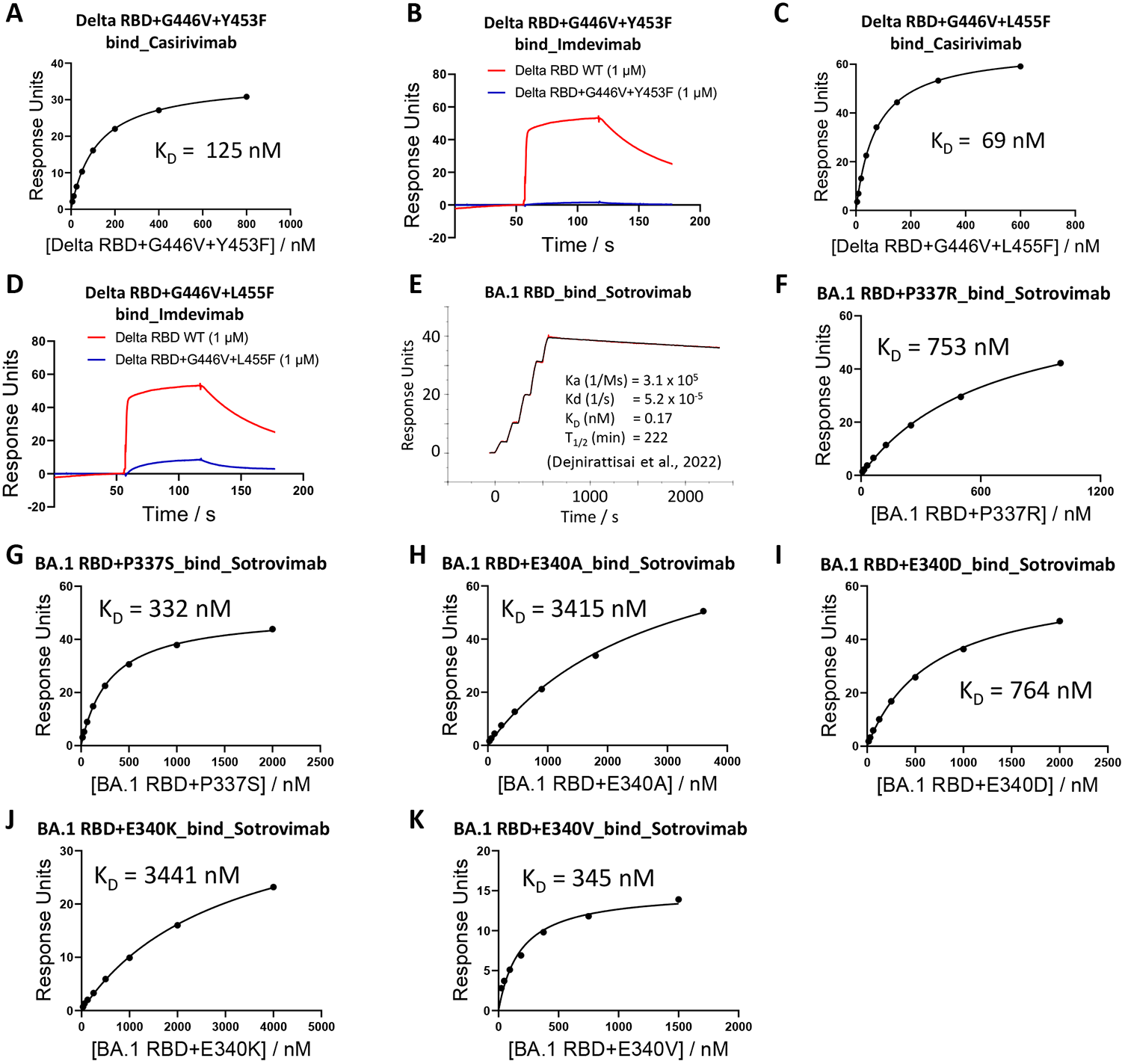
Surface plasmon resonance (SPR) analysis of interaction between Delta and BA.1 RBD mutants and therapeutic mAbs. (A, C) 1:1 binding equilibrium analysis of binding of Delta RBD mutants to casirivimab, with affinity values shown. (B, D) Binding of Delta RBD+G446V+Y453F and Delta RBD+G446V+L455F to imdevimab is severely reduced compared to that of wild-type Delta RBD, so that the binding could not be accurately determined, as shown by a single-injection of 1 μM RBD over sample flow cells containing imdevimab. (E) Sensorgram showing the binding of wild-type BA.1 RBD to sotrovimab, with affinity and kinetic parameters shown (published in Dejnirattisai et al., 2022). (F-K) 1:1 binding equilibrium analysis of binding of BA.1 RBD mutants to sotrovimab, with affinity values shown. Related to **Figure S2** and **Table 3**.

**Table S1.** Combinations of SARS-CoV-2 variants, gene sequences and treatments examined for resistance mutations.

| Treatment | Genes | Variants |
| --- | --- | --- |
| Sotrovimab | Spike | Delta, BA.1, BA.2 |
| Casirivimab and<br>imdevimab | Spike | Delta, BA.1, BA.2 |
| Remdesivir | NSP7, NSP8, NSP9, NSP10, NSP12 | Delta, BA.1, BA.2 |
| Molnupiravir | NSP7, NSP8, NSP9, NSP10, NSP12 | Delta, BA.1, BA.2 |
| Paxlovid | NSP5 | Delta, BA.1, BA.2 |

## STAR Methods

### Resource availability Lead Contact

Resources, reagents and further information requirement should be forwarded to and will be responded to by the Lead Contact, David Stuart

### Materials Availability

Reagents generated in this study are available from the Lead Contact with a completed Materials Transfer Agreement.

### Study population

In April 2020, the UK established a national program of SARS-CoV-2 genomic surveillance through which viruses from a random sample of population positives in the community and hospital have been routinely sequenced (, 2020). In addition, a protocol was introduced to enhance sequencing coverage of those receiving treatment in hospitals and within the community (including pre-treatment and follow-up sampling). Patients on treatment were linked to their genetic sequences through their COG-IDs. The present analysis includes all patients who have received treatment in the UK, for whom at least one sample had been collected by 12 April 2022 and for whom a viral genetic sequence was available.

At this date, the five therapeutic interventions deployed across the population included casirivimab/imdevimab, sotrovimab, molnupiravir, remdesivir and paxlovid (nirmatrelvir plus ritonavir). For each patient, data available include: date of sample, therapeutic intervention/ treatment and date of treatment.

Surveillance of coronavirus disease 2019 (Covid-19) testing and vaccination is undertaken under Regulation 3 of the Health Service (Control of Patient Information) Regulations 2002 to collect confidential patient information (www.legislation.gov.uk/uksi/2002/1438/regulation/3/made. opens in new tab) under Sections 3(i) (a) to (c), 3(i)(d) (i) and (ii), and 3. The genomic surveillance study protocol (https://www.gov.uk/government/publications/covid-19-genomic-surveillance-of-patients-who-are-treated-with-neutralising-monoclonal-antibody-or-immunosuppressed) was subject to an internal review by the UKHSA England Research Ethics and Governance Group and was found to be fully compliant with all regulatory requirements. Given that no regulatory issues were identified, and that ethics review is not a requirement for this type of work, it was decided that a full ethics review would not be necessary.

### Sequence datasets

The pipeline used to collect and process raw SARS-CoV-2 sequence data and sample-associated metadata across the UK genomic surveillance network has been previously described (Nicholls et al., 2021). The ARCTIC protocol was employed to amplify SARS-CoV-2 samples (Lambisia et al., 2022). Sequencing platforms included Illumina and Oxford Nanopore Technologies. Sequences were aligned to the reference SARS-CoV-2 genome (NCBI NC_045512.2). COVID lineages were assigned using Pango (O’Toole et al., 2022).

### Analysis of pre- and post-treatment sequences

All sequences from patients known to have undergone treatment were downloaded from CLIMB. Genome alignments were split into gene regions (spike, NSP5, NSP7, NSP8, NSP9, NSP10, NSP12 and NSP14) and translated to amino acids for analysis. Analyses were conducted for each treatment on the proteins they are theorised to interact with. Analyses were split by variant (Delta, Omicron BA.1 and Omicron BA.2), with Delta sublineages (B.1617.2 and all AY lineages) all classified as Delta. As such, each analysis was conducted independently on every treatment, variant and gene region combination of interest (**Table S1**).

Pre-treatment sequences are those obtained from patients with a sequenced sample within one week prior to treatment initiation (including the day of treatment initiation). The analysis was repeated with a range of cut-offs for defining post-treatment sequences, including post-treatment sequences only if they were sampled at least 1,5,10, or 14 days after treatment. Our main analysis uses the 10-day cut-off, and 1,5 and 14 days are presented as a sensitivity analysis. For each analysis, we split the dataset into pre-and post-treatment sequences. At each site in the alignment, the amino acid frequency was calculated in pre- vs post-treatment sequences, and Fisher’s exact test was used to determine whether this probability distribution diverged from the null expectation. In this way, sites that display unexpected differences in amino acid frequencies were identified, and the specific amino acid changes highlighted. Analyses were conducted at the patient-level rather than at the sequence level, so that if a patient had multiple pre- or post-treatment sequences a single sequence was retained, with sequences diverging from the wild-type favoured.

### Analysis of UK genomic database

All Delta (n=763,511), BA.1 (n=742,992) and BA.2 (n=XX) sequences from September 2021 onwards were downloaded from CLIMB, translated to amino acids and split into proteins using an in-house script. For each amino acid site identified in our analysis, amino acid frequencies were tabulated and calculated as proportions of the total number of sequences with a readable amino acid.

### Structural modelling/Mapping of drug interaction sites

Structural models of RBD-nmAbs complexes were generated by superposition of PDB:7ORB (RBD with L452R) and Omicron RBD (PDB:7TLY) with complexes of RBD-casirivimab/imdevimab (PDB:6XDG) and RBD-sotrovimab (PDB:7BEP) respectively, using program SHP (Stuart et al., 1979) to align the RBD domains. Models of PDB:7ORB RBD docked with casirivimab/imdevimab and Omicron RBD docked with sotrovimab were extracted and analysed at drug interaction sites using Coot (Casanal et al., 2020). Molecular graphics images were generated using UCSF ChimeraX (Pettersen et al., 2021).

### Sera from Pfizer vaccinees

Pfizer vaccine serum was obtained from volunteers who had received three doses of the BNT162b2 vaccine (Pfizer/BioNTech). Vaccinees were Health Care Workers, based at Oxford University Hospitals NHS Foundation Trust, not known to have prior infection with SARS-CoV-2 and were enrolled in the OPTIC Study as part of the Oxford Translational Gastrointestinal Unit GI Biobank Study 16/YH/0247 [research ethics committee (REC) at Yorkshire & The Humber – Sheffield] which has been amended for this purpose on 8 June 2020. The study was conducted according to the principles of the Declaration of Helsinki (2008) and the International Conference on Harmonization (ICH) Good Clinical Practice (GCP) guidelines. Written informed consent was obtained for all participants enrolled in the study. Participants were sampled approximately 28 days (median 31, range 28-56), after receiving a third “booster” dose of Pfizer/BioNtech BNT162b2 mRNA Vaccine, 30 micrograms, administered intramuscularly after dilution (0.3 mL each), 17-28 days apart for dose 1 and 2, then approximately 9 months apart (range 253-300) for dose 2 and 3. The mean age of vaccinees was 42 years (range 30-59), 10 male and 9 female.

### Plasmid construction and pseudotyped lentiviral particles production

Pseudotyped lentivirus expressing SARS-CoV-2 S proteins for ancestral strains (Victoria, Delta and BA.1) were constructed as described before (Nie et al., 2020; Liu et al., 2021, Nutalai et al., 2022) with some modifications. A similar strategy was applied for all variant constructs. Delta and BA.1 were used as the template and the constructs were cloned by PCR amplification of vector and inserts, followed by Gibson assembly. To generate the insert fragments, the overlapping primers for all individual variants were used separately to amplify, together with two primers of pcDNA3.1 vector (pcDNA3.1_BamHI_F and pcDNA3.1_Tag_S_EcoRI_R). The pcDNA3.1 vector was also amplified using pcDNA3.1_Tag_S_EcoRI_F and pcDNA3.1_BamHI_R primers. The primer pairs used in this study are shown in supplementary (**Table S1**). All constructs were verified by Sanger sequencing after plasmid isolation using QIAGEN Miniprep kit (QIAGEN). The resulting S gene-carrying pcDNA3.1 was used for generating pseudoviral particles together with the lentiviral packaging vector and transfer vector encoding luciferase reporter.

### Pseudoviral neutralization assay

The details of pseudoviral neutralization test were described previously (Liu et al., 2022; Nie et al., 2020) with some modifications. Briefly, four-fold serial dilution of each mAb was incubated with pseudoviral particles at 37 °C, 5% CO_2_ for 1 h. The stable HEK293T/17 cells expressing human ACE2 were then added to the mixture at 1.5 × 10^4^ cells/well. At 48 h post transduction, culture supernatants were removed and 50 µL of 1:2 Bright-GloTM Luciferase assay system (Promega, USA) in 1x PBS was added into each well. The reaction was incubated at room temperature for 5 min and the firefly luciferase activity was measured using CLARIOstar (BMG Labtech, Ortenberg, Germany). The percentage of neutralization was calculated relative to the control. Probit analysis was used to estimate the value of dilution that inhibits half of the maximum pseudotyped lentivirus infection (PVNT50). To determine the neutralizing activity of vaccine sera, 3-fold serial dilutions of samples were incubated with pseudoviral particles for 1 hr and the same strategy as mAb was applied. The primer sequences used to generate pseudoviruses are listed in **Table S2**.

### Cloning of RBDs

To generate the His-tagged construct of RBDs, site-directed PCR mutagenesis was performed using the Delta or BA.1 pseudovirus plasmid construct as the template, or pseudovirus plasmid construct containing the desired RBD mutant was used as the template for amplification of the RBD gene fragment.

The template, primers and expression vectors used for cloning of each RBD are shown in **Table S3 and t**he primer sequences are shown in **Table S4**.

Cloning was performed using the ClonExpress II One Step Cloning Kit (Vazyme). The Constructs were verified by Sanger sequencing after plasmid isolation using QIAGEN Miniprep kit (QIAGEN).

### Production of RBDs

Plasmids encoding RBDs were transfected into Expi293F™ Cells (ThermoFisher) by PEI, cultured in FreeStyle™ 293 Expression Medium (ThermoFisher) at 30 °C with 8% CO_2_ for 3 days. The conditioned medium was diluted 1:2 into binding buffer (50 mM sodium phosphate, 500 mM sodium chloride, pH 8.0). RBDs were purified with a 5 mL HisTrap nickel column (GE Healthcare) through His-tag binding, followed by a Superdex 75 10/300 GL gel filtration column (GE Healthcare) in 10 mM HEPES and 150 mM sodium chloride.

### Surface Plasmon Resonance

The surface plasmon resonance experiments were performed using a Biacore T200 (GE Healthcare). All assays were performed with a running buffer of HBS-EP (Cytiva) at 25⍰°C. A Protein A sensor chip (Cytiva) was used. The mAb as indicated was immobilised onto the sample flow cell of the sensor chip. The reference flow cell was left blank.

To determine the binding kinetics, RBD was injected over the two flow cells at a range of five concentrations prepared by serial two-fold dilutions, at a flow rate of 30⍰μ⍰min^−1^ using a single-cycle kinetics programme. Running buffer was also injected using the same programme for background subtraction. All data were fitted to a 1:1 binding model using Biacore T200 Evaluation Software 3.1. To determine the binding affinity (where kinetics were difficult to determine), RBD was injected over the two flow cells at a range of concentrations prepared by serial two-fold dilutions, at a flow rate of 30⍰μ⍰min^−1^. Running buffer was also injected using the same programme for background subtraction. All KD data were fitted to a 1:1 binding model using Biacore T200 Evaluation Software 3.1; the figures were plotted with GraphPad Prism 9.

To compare the binding profiles between Delta RBD+G446V / Delta RBD+G446V+Y453F / Delta RBD+G446V+L455F and Delta RBD WT for imdevimab, a single injection of RBD was performed over the two flow cells at 1 μM, at a flow rate of 30⍰μ⍰min^−1^. Running buffer was also injected using the same programme for background subtraction. The sensorgrams were plotted using Prism9 (GraphPad).

